# U1 snRNP regulates chromatin retention of noncoding RNAs

**DOI:** 10.1101/310433

**Authors:** Yafei Yin, J. Yuyang Lu, Xuechun Zhang, Wen Shao, Yanhui Xu, Pan Li, Yantao Hong, Qiangfeng Cliff Zhang, Xiaohua Shen

**Affiliations:** Tsinghua-Peking Center for Life Sciences, School of Medicine and School of Life Sciences, Tsinghua University, Beijing 100084, China; MOE Key Laboratory of Bioinformatics, Beijing Advanced Innovation Center for Structural Biology, Center for Synthetic and Systems Biology, School of Life Sciences, Tsinghua University, Beijing 100084, China

**Keywords:** RNA-chromatin localization, U1 snRNA, lncRNA, PROMPTs and eRNA, splicing

## Abstract

Thousands of noncoding transcripts exist in mammalian genomes, and they preferentially localize to chromatin. Here, to identify *cis*-regulatory elements that control RNA-chromatin association, we developed a high-throughput method named RNA element for subcellular localization by sequencing (REL-seq). Coupling REL-seq with random mutagenesis (mutREL-seq), we discovered a key 7-nt U1 recognition motif in chromatin-enriched RNA elements. Reporter assays indicated a direct role for U1 snRNP recognition in regulating RNA-chromatin localization. Globally, U1 motifs and U1 binding are strongly enriched in long noncoding RNA (lncRNA) transcripts. Inhibition of U1 snRNA, and of U2 to a lesser degree, led to global reduction in chromatin association of hundreds of lncRNAs. For promoter- and enhancer-associated noncoding RNAs, U1 binds to their genomic neighborhoods, and their chromatin association depends on both U1 and U2 snRNAs. These findings reveal that U1 snRNP, perhaps together with the splicing machinery, acts widely to promote the chromatin association of noncoding transcripts.

## Introduction

The subcellular localization of RNA often correlates with its function and how it is regulated. Most protein-coding transcripts are exported to the cytoplasm for protein synthesis (Kohler and Hurt, 2007), whereas some intron-retaining transcripts are retained in the nucleus to reduce the levels of host transcripts that are not essential for the cell (Boutz et al., 2015; Braunschweig et al., 2014). Pervasive transcription in mammalian genomes produces hundreds of thousands of noncoding RNA (ncRNA) transcripts (Djebali et al., 2012). It has been proposed that the human genome expresses ~19,175 potentially functional long ncRNAs (lncRNAs) (Hon et al., 2017). In addition, active promoters and enhancers often generate sense and antisense unstable transcripts named promoter upstream transcripts (PROMPTs, or transcription start site-associated RNAs – TSSa-RNAs) and enhancer-associated RNAs (eRNAs), respectively (Core et al., 2008; De Santa et al., 2010; Kim et al., 2010; Preker et al., 2011; Preker et al., 2008; Seila et al., 2008). Remarkably, both lncRNAs and the class of promoter- and enhancer-associated ncRNAs are preferentially located on chromatin in the nucleus of the cell (Cabili et al., 2015; Derrien et al., 2012; Li et al., 2016; Schlackow et al., 2017).

While some *trans-acting* lncRNAs like *Malat1* and *Neat1* bind to distant genomic regions on chromatin, many other cis-acting lncRNAs, including *Xist, RepA, Tsix, Coolair, Airn, Kcnq1ot1, Haunt* and *Evx1as*, bind to their own transcription sites and chromatin neighborhoods to fine-tune gene expression and chromatin structure (Chen, 2016; Chu et al., 2011; Latos et al., 2012; Lee and Bartolomei, 2013; Luo et al., 2016; Pandey et al., 2008; Swiezewski et al., 2009; Yan et al., 2017; Yin et al., 2015). For example, *Malat1*, a conserved and abundant nuclear lncRNA which often shows altered expression in human cancer, is mainly localized in the nuclear speckles and binds to thousands of genomic DNA sites *in trans* to regulate transcription, pre-mRNA splicing and nuclear architecture (Arun et al., 2016; Engreitz et al., 2014; Tripathi et al., 2010; West et al., 2014). The X-inactive specific transcript (*Xist*) is a classic *cis*-acting, chromatin-associated lncRNA involved in X chromosome inactivation (XCI) in mammals (Payer and Lee, 2008). *Xist* expression and accumulation in *cis* on the prospective inactive X chromosome recruits epigenetic repressors and triggers silencing of the whole X chromosome (Jegu et al., 2017; Lessing et al., 2013; Plath et al., 2002). Despite the well-reported phenomenon of chromatin association of lncRNAs, the mechanisms that regulate RNA chromatin tethering remain elusive.

Cytoplasmic mRNAs and chromatin-retained lncRNA transcripts utilize RNA polymerase II and a similar set of RNA processing factors for their synthesis, packaging, processing, and turnover (Derrien et al., 2012). However, they may diverge at some point along their maturation pathway before reaching their final destination. A recent study in yeast reported that polymerase-associated factor 1 (PAF1) controls the nuclear export of mRNAs and preferential nuclear retention of the majority of lncRNAs and some mRNAs (Fischl et al., 2017). Proteome-wide surveys for proteins that bind to mRNAs and lncRNA transcripts have suggested that 3’ cleavage/polyadenylation factors may restrict nuclear export of lncRNAs and pre-mRNAs by modulating splicing and nuclear RNA surveillance pathways (Baejen et al., 2014; Tuck and Tollervey, 2013). Analysis of nascent transcription across the human genome has shown that co-transcriptional splicing of most lncRNAs, unlike protein-coding genes, is inefficient and lncRNAs often exhibit frequent transcription termination at multiple positions within the gene body (Schlackow et al., 2017; Tilgner et al., 2012). Quantitative and qualitative differences in lncRNA and pre-mRNA processing may contribute in part to the nuclear retention of some short-lived lncRNA transcripts (Schlackow et al., 2017). Indeed, many lncRNAs and ncRNA transcripts exhibit fast turnover and are prone to exosome-mediated RNA degradation in the nucleus (Andersson et al., 2014; Lubas et al., 2015; Pefanis et al., 2015). However, stable lncRNAs like *Xist* and *Malat1* may employ mechanisms beyond nuclear surveillance for their chromatin tethering.

RNA splicing is performed by a spliceosome that assembles on each intron and predominantly comprises small nuclear ribonucleoproteins (snRNPs), including U1, U2, U4, and U5, and U6, snRNPs in equal stoichiometry (Kaida et al., 2010; Lerner et al., 1980). U1 snRNP plays an essential role in defining the 5’ splice site by RNA:RNA base pairing via the 9-nucleotide (nt) sequence at the 5’ end of U1 snRNA (Will and Luhrmann, 2011). Independent from its role in splicing, U1 snRNP protects pre-mRNAs from premature cleavage and polyadenylation at cryptic polyadenylation signals (PASs) in introns, a process termed telescripting (Berg et al., 2012). Despite a handful of studies on nuclear retention of intron-retaining mRNA transcripts, the role of U1 snRNP recognition and binding in retaining noncoding RNAs on chromatin remains unclear.

Evidence shows that intrinsic mRNA elements, like the constitutive transport element (CTE) and the RNA transport element (RTE), serve as the “postage” for RNA export and cytoplasmic localization of some viral RNAs and a small subset of mRNAs (Hammarskjold, 2001; Nappi et al., 2001). Deletion analysis of the lncRNAs *Xist, Malat1* and *BORG* identified large stretches of RNA sequences that contribute to their nuclear localization (Miyagawa et al., 2012; Ridings-Figueroa et al., 2017; Sunwoo et al., 2017; Wutz et al., 2002; Zhang et al., 2014). We conjecture that cytoplasm-localized mRNAs and chromatin-retained ncRNA transcripts may differ at the sequence level so that distinct *trans-acting* protein factors can act upon them. The fate of an RNA transcript, in terms of its subcellular localization, may be determined by intrinsic sequence motifs.

To test this hypothesis and to reveal the cis-regulatory RNA elements that control RNA localization, we developed high-throughput screening methods named REL-seq and mutREL-seq. These screens led us to uncover a 7-nt U1 recognition motif that regulates RNA chromatin localization. Globally, lncRNAs as a class show strong and significant enrichment of U1 recognition motifs and U1 snRNP binding compared to cytoplasm-enriched mRNA. Importantly, inhibition of U1 snRNP, and of U2 to a lesser degree, caused a global reduction in chromatin associations of hundreds of lncRNAs. In addition, the chromatin tethering of PROMPTs and eRNAs relies on both U1 and U2 snRNAs. These results reveal U1 recognition as a novel cis-regulatory signal for RNA-chromatin association, and demonstrate a prevalent involvement of U1 snRNP, and perhaps the splicing machinery, in retaining noncoding RNA transcripts on the chromatin.

## Results

### Strategy for REL-seq

We fractionated chromatin-bound RNA in mouse embryonic stem cells (mESCs) and compared RNA-seq profiles of chromatin RNA to total RNA in the cell. We also analyzed a previously published RNA-seq dataset of chromatin and total RNA in human K562 cells (Tilgner et al., 2012). Consistent with previous reports (Derrien et al., 2012; Schlackow et al., 2017), lncRNAs as a class are significantly enriched in the chromatin fraction in both mouse and human cells (Figure 1A). We sought to perform an unbiased screen to comprehensively identify intrinsic *cis*-regulatory signals embedded within an RNA transcript that contribute to its chromatin association.

**Figure 1.**
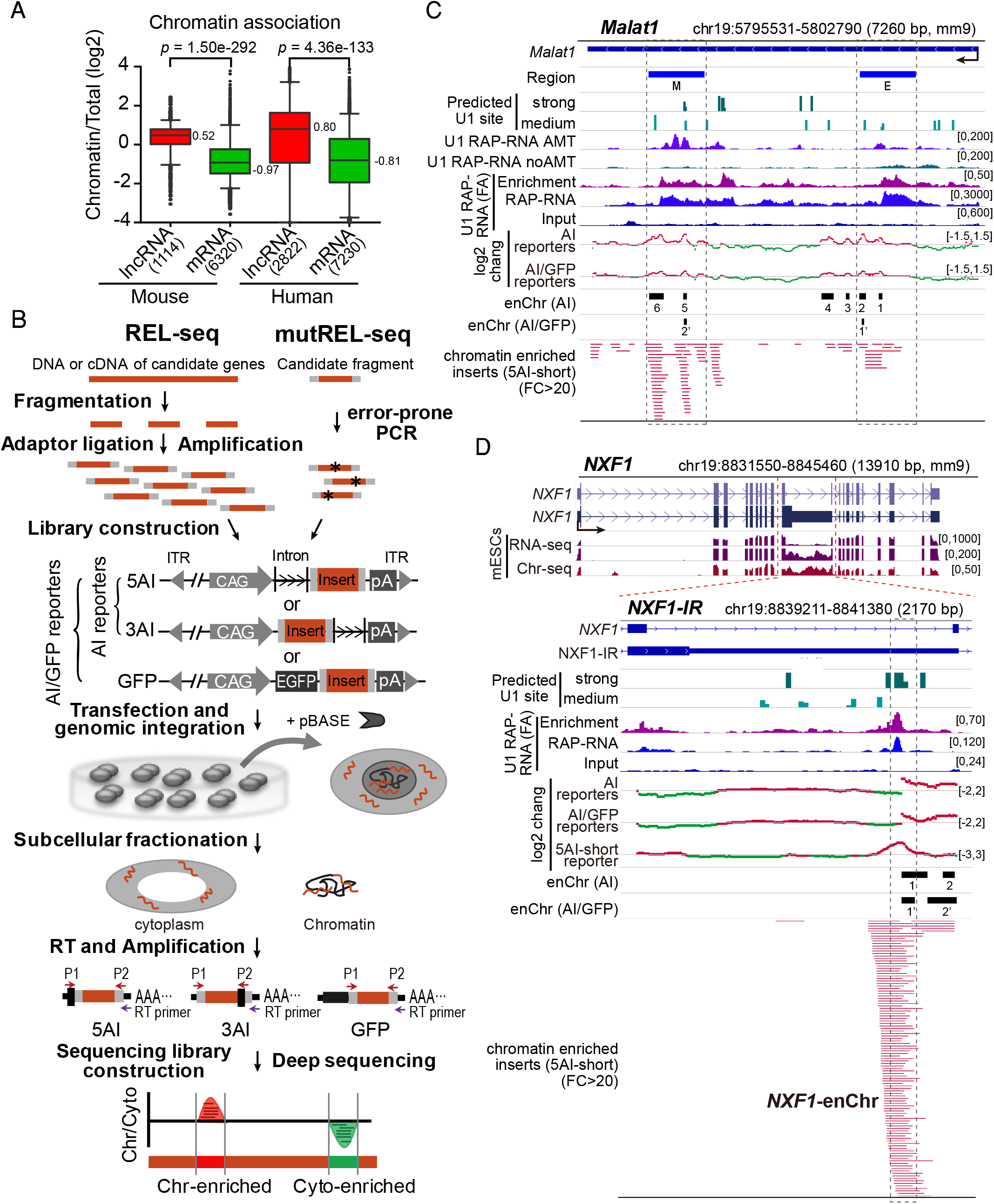
REL-seq identifies cis-elements that contribute to RNA subcellular localization. (A) Chromatin enrichment comparison between lncRNAs and coding genes in human and mouse. All lncRNAs with FPKM > 1 and all coding genes with FPKM > 5 were used for the comparison. T-tests were used to compare the data for two indicated samples. The median value for each sample is labeled beside the box-plot. (B) Pipeline for REL-seq and mutREL-seq. Candidate DNAs or double-strand cDNAs were fragmented, ligated with in-house-designed adaptors (Table S5), and amplified. The PCR products were further inserted into 5AI, 3AI, or GFP reporter vectors, which are based on the *PiggyBac* transposon system. ITR: inverted terminal repeat sequences of the *PiggyBac* transposon system. After the plasmids were transfected into mESC or 293T cells (co-transfected with plasmids expressing the pBASE transposase) and integrated into the genomic DNA, a subcellular fractionation assay was performed to separate the cells into cytoplasmic, nucleoplasmic and chromatin fractions. RNAs from different fractions were extracted, reverse-transcribed (RT), and amplified with specific primers (purple arrows: RT primers for reverse-transcription, red arrows: P1, P2 primers for amplification; see Table S5). Libraries for deep sequencing were further constructed. Sequences that were enriched in different subcellular fractions were identified by comparing the read intensities or insert abundance in the chromatin fraction with that of the cytoplasmic fraction. Left brackets highlight the reporters that were included in the AI/GFP screen or the AI screen. For mutREL-seq, candidate fragments were randomly mutagenized through error-prone PCR, and the products were further inserted into reporter vectors and subjected to downstream procedures similar to those described for REL-seq. Asterisks represent the mutation sites. (C) REL-seq identifies enChrs in the mouse *Malat1* transcript. The locations of the mouse homologous regions E and M in human *Malat1* are shown as thick blue lines. In addition to the RNA-seq signal of U1 RAP-RNA with formaldehyde crosslinking (U1 RAP-RNA FA), RNA-seq signal of U1 RAP-RNA with AMT crosslinking (U1 RAP-RNA AMT) and respective control (U1 RAP-RNA noAMT) are also shown. The U1 RAP-RNA AMT used 4’-aminomethyltrioxalen (AMT) to crosslink the cells, which generates inter-strand crosslinks between uridine bases through with, detect the direct RNA-RNA interactions of highly expressed transcripts (Engreitz et al., 2014). The chromatin-enriched inserts from the 5AI-short reporter screen are shown at the bottom (fold enrichment > 10). The dashed grey rectangles highlight the read density information and identified enChrs located in the mouse homologous regions E and M. (D) REL-seq identifies enChrs in the mouse *NXF1-IR* region. The average fold changes of inserts in the AI/GFP reporters, AI reporters and the 5AI-short reporter are shown. The chromatin-enriched inserts of the 5AI-short reporter (fold enrichment > 20) are shown at the bottom. The dashed grey rectangle highlights the *NXF1*-enChr region.

The strategy is to express a random RNA fragment alone or fuse it with a GFP mini-gene, and then analyze its subcellular location. Tens of thousands of RNA fragments can be inserted and expressed in reporter plasmids, and RNAs that are specifically localized in a particular compartment can be identified by high-throughput sequencing. The underlying sequence of an RNA that is enriched in a particular subcellular compartment may contain a *cis* signal for its localization. We named this method as RNA elements for subcellular localization by sequencing (REL-seq) (Figure 1B).

To generate REL-seq libraries, cDNA or genomic sequences from candidate genes were randomly fragmented and inserted into two types of reporter vector. One type, herein referred to as the AI reporter, contains only an *ACTB* intron, and was used to distinguish spliced mature RNAs from nascent transcripts. Random DNA fragments were inserted either upstream (3’) or downstream (5’) of the *ACTB* intron, and designated as 3AI and 5AI, respectively. Mature RNAs expressed in AI reporters are solely derived from the inserted DNA fragments and the subcellular localization of the RNA therefore depends on the inserted DNA sequences. In the second type of reporter, a random RNA sequence is fused to a *GFP* mini-gene to screen for strong chromatin retention sequences that can overcome the nuclear export effect of *GFP* (Figure 1B). We reasoned that the protein-coding open reading frame of *GFP* RNA would likely promote its cytosolic transport for translation. RNA fragments that show enriched chromatin localization in both AI and GFP (AI/GFP) reporter screens may represent stronger and higher-confidence signals for chromatin association.

We performed REL-seq for 9 representative transcripts, including five chromatin-retained lncRNA transcripts *(Malat1, Neat1, Xist, NR_028425,* and a portion of *Tsix* that is reverse complementary to *Xist* RNA), one chromatin-retained mRNA *(NXF1),* and three cytoplasmic mRNAs *(NCL, NANOG* and *ACTB)* (Figures S1A-S1C). *Malat1, Neat1, Xist,* and *Tsix* are conserved, well-characterized lncRNAs with predominantly nuclear and chromatin localization in both human and mouse. *NR_028425* is a mouse-specific, uncharacterized lncRNA and is strongly enriched on chromatin in mESCs (Figure S1A). *NXF1* (nuclear RNA export factor 1) mediates nuclear export of most mRNAs (Natalizio and Wente, 2013). On the other hand, *NXF1* expression is regulated by the nuclear retention of an alternatively spliced *NXF1* mRNA, which retains intron 10, designated as *NXF1-IR* (Li et al., 2006; Sasaki et al., 2005). As *NXF1-IR,* but not normally spliced *NXF1* mRNA, is strongly enriched on chromatin in mESCs (Figures 1D and S1A), we speculated that strong RNA signals for nuclear retention may exist within this 2-kb intron of *NXF1-IR.*

In order to balance signal resolution and the length of fragmentation without disrupting regulatory RNA sequences or secondary structures required for nuclear retention, we prepared REL-seq libraries with DNA inserts ranging in length from 150 to 600 bp (Figures S1D, S1E, and Table S1). To minimize cell-specific effects and identify RNA signal sequences conserved in human and mouse, we constructed a total of 8 REL-seq libraries in both mESCs and human HEK 293T cells. These libraries harbor 462,885 unique RNA inserts covering the 60,358-nt candidate RNAs for ~395 times on average per library (Table S1), indicating saturated coverage of fragments for the 9 RNA transcripts analyzed. As DNA can be inserted in both sense and reverse orientations during REL-seq library construction, nearly half of the inserts were expected to express the reverse complement sequences of the candidate RNAs, which thus serve as internal controls to assess the specificity of the sense RNA signals identified.

### REL-seq screens effectively identify chromatin-enriched RNAs (enChrs)

To identify chromatin-enriched RNA fragments (enChrs), we used a stringent cutoff to select fragments that show significantly higher read densities in the chromatin (chr) fraction than in the cytoplasmic (cyto) fraction (average chr/cyto ratio >1.5 and *p* < 0.05, t-test of all samples from seven AI screens or from eleven screens using AI and GFP reporters). In total, we identified 26 and 14 enChrs by AI or AI/GFP reporter screens, respectively, in the sense orientation of the host chromatin-associated RNAs but not in cytoplasm-located mRNAs (Figures 1C, 1D, S1F-S1H, and S2, and Table S2). Few enChrs (5 in AI and none in AI/GFP screens) were in the reverse complementary direction (Table S2). This result is consistent with host RNA subcellular localization, further supporting the notion that intrinsic signals lie within the RNA transcripts to instruct their subcellular localization. Additionally, we identified several enriched cytoplasmic RNA regions in mRNA candidates (Figure S1F and Table S2). Together, these results indicated the effectiveness of the REL-seq strategy to identify cis-regulatory RNA sequences involved in the chromatin localization of RNA. Because very stringent selection criteria were employed across all REL-seq datasets generated using different libraries in three AI and GFP reporter vectors in both human and mouse cell lines, the enChrs we identified may underrepresent the total number of regulatory RNA signals for chromatin association in the cell.

We compared the enChrs with previously reported chromatin-localized regions. *Malat1* harbors six AI enChrs and two AI/GFP enChrs. Most of them are located within two 1 -kb regions, homologous to regions E and M in human *Malat1* (Figure 1C). These two regions were previously reported to be critical for human *Malat1* localization to nuclear speckles (Miyagawa et al., 2012). The *Xist* RNA harbors eight AI enChrs and five AI/GFP enChrs (Figure S2A), some of which are located within or near to the repeat regions A, D, E and F (Chen et al., 2016; Lv et al., 2016; Ridings-Figueroa et al., 2017; Sunwoo et al., 2017; Wutz et al., 2002). For example, one strong enChr was identified at the 5’ end of *Xist* overlapping with the repeat A, which is known to be important for XCI (Chu et al., 2015; McHugh et al., 2015; Wutz et al., 2002). Deletion of either repeat D or F impairs XCI (Chen et al., 2016; Lv et al., 2016; Wutz et al., 2002); however, the mechanistic link to their function in XCI is unclear. Identification of several enChrs within repeats D and F suggests that these repeats may regulate XCI by promoting *Xist* chromatin association. In addition, multiple enChrs are located in the last exon of *Xist,* which contains repeat E. This finding is consistent with the reports that successive truncations of repeat E along with its downstream sequences attenuate chromatin binding of *Xist* and impair XCI (Ridings-Figueroa et al., 2017; Sunwoo et al., 2017). Thus, REL-seq analysis of *Xist* and *Malat1* not only identified all known regions that regulate their chromatin association, but further narrowed these regions down to relatively short sequences. Multiple enChrs along both transcripts may act synergistically to enhance their chromatin binding.

### MutREL-seq reveals a U1 motif critical for the chromatin binding of NXF1-enChr

REL-seq identified a pool of chromatin-enriched RNA sequences, mainly ranging between 50~500 nt in length depending on the sizes of the random DNA fragments inserted into the REL-seq libraries (Table S2). In order to further increase the resolution of the analysis and identify key motifs or residues that contribute to RNA chromatin localization, we sought to modify the REL-seq strategy by combining it with random mutagenesis (mutREL-seq, Figure 1B).

REL-seq revealed two strong enChrs that are located towards the 3’ end of the retained intron of *NXF1-IR* (Figure 1D). In the #1 enChr site of *NXF1-IR,* 116 unique chromatin-enriched RNA fragments (chr/cyto ratio > 20-fold) were uncovered in one REL-seq screen using the 5AI reporter (Figure 1D). Because of its remarkable enrichment on chromatin, this 162-nt region, designated as NXF1-enChr (highlighted by the grey dashed box in Figure 1D), was subjected to mutagenesis by error-prone PCR. Mutations that disrupt or promote chromatin binding of NXF1-enChr were identified by REL-seq. Among a total of 469 mutation events (Figures S3A-S3D), 23 RNA mutants showed impaired chromatin binding of NXF1-enChr RNA with a > 2-fold increase of the cytoplasm to chromatin (cyto/chr) ratio. In contrast, very few mutations (only 3) exhibited enhanced chromatin binding (cyto/chr < −1.25-fold), consistent with the role of NXF1-enChr RNA as a strong signal to promote RNA-chromatin association (Figure 2A). Remarkably, 19 out of 23 mutations (cyto/chr > 2-fold) are located within a 7-nt region spanning from nucleotide position 39 to 45 of the 162-nt *NXF1*-enChr RNA (Figure 2A).

**Figure 2.**
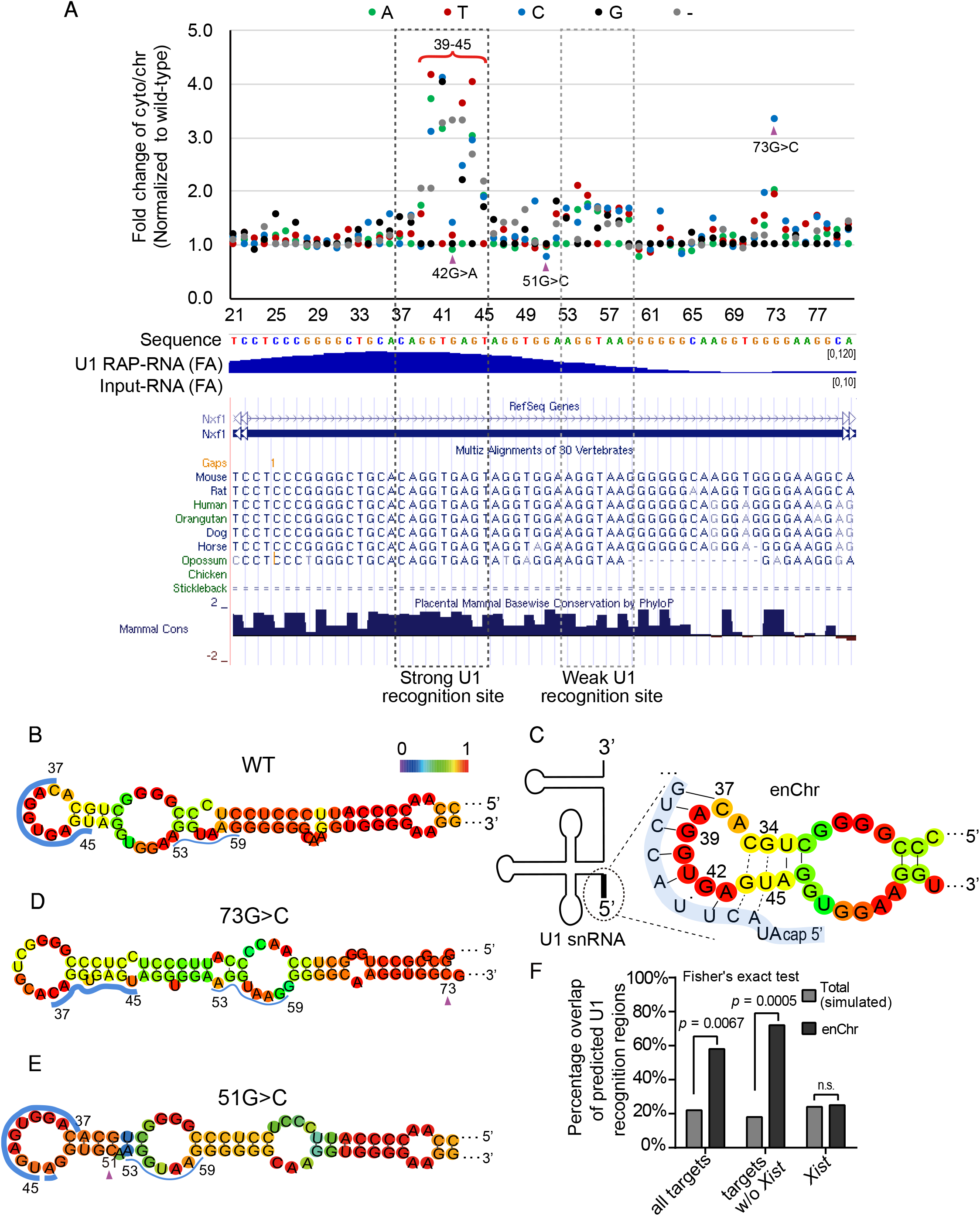
MutREL-seq reveals a U1 motif critical for the chromatin binding of NXF1-enChr. (A) MutREL-seq identifies a conserved motif that contributes to nuclear retention of NXF1-enChr. Upper panel, fold changes of the cytoplasmic fraction relative to the chromatin fraction of different mutations at positions 21-80 of the NXF1-enChr region (fold changes of different mutations are normalized to wild-type). Middle, RNA-seq signals of U1 RAP-RNA with formaldehyde crosslinking (U1 RAP-RNA FA) and the respective input (Input-RNA FA) are shown. Bottom, the conservation in vertebrates of each position in the upper panel of the NXF1-enChr region is shown. The U1 snRNP recognition site and a weak U1 recognition site are highlighted in the dashed black rectangles. The red bracket at the top highlights positions 39-45. The three purple triangles highlight the 42G>A, 51G>C, and 73G>C mutations. (B) Predicted secondary structure of the NXF1-enChr region (partial). The U1 snRNP targeting region at position 37-45 is highlighted with a thick blue line, and weak U1 recognition site is highlighted with thin blue line. The colored scale bar in Figures 2B-E represents the probability of base-pairing or being unpaired (red, high probability; blue, low probability). (C) Predicted secondary structure of the RNA interaction between the NXF1-enChr region and U1 snRNA. The schematic diagram of U1 snRNA is shown on the left side, and the intron-binding site at the 5’ end is highlighted with a thick black line. The zoomed-in figure shows the sequence of the 5’ region of U1 snRNA, which is highlighted by a thick transparent blue line. The base-pairing between U1 snRNA and NXF1-enChr is shown. The U:G wobble base-pairing at position 42 of *NXF1-*enChr is indicated by a dot. For positions 44 and 45 of NXF1-enChr, dashed lines are used to show the base-pairing with U1 snRNA, as these bases may also pair with positions 34 and 35 of *NXF1-* enChr. (D, E) Predicted secondary structure of *NXF1-enChr* RNA with the 73G>C (D) and 51G>C (E) mutations. The purple triangles highlight the mutation site. (F) The percentage overlap of predicted U1 snRNP recognition sites with simulated total regions or enChrs. The p-value of Fisher’s exact test is shown at the top of each pair of compared samples.

The 7-nt core motif and nearby sequences of NXF1-enChr are highly conserved in eukaryotes (Figure 2A). RNA secondary structure prediction analysis revealed that the NXF1-enChr RNA forms a stable secondary structure containing four protruding loops (free energy of the thermodynamic ensemble: −69.49 kcal/mol) (Figures S3E and S3F). Interestingly, 5 of the core 7 nucleotides reside within an 8-nt protruding loop (marked by a thick blue line in Figures 2B and S3E). The 7-nt sequence along with two upstream nucleotides (5’ CA) comprises a U1 recognition site (5’ CAGGUGAGU), and base pairs nearly perfectly with the 5’ targeting sequence of U1 snRNP (5’ACUUACCUG) (Figure 2C). The underlines in the above sequences indicate the G:U wobble base-paired nucleotide.

The decreased chromatin association observed in 19 mutations in each of the 7-nt bases likely results from their disrupted base-pairing with the targeting sequence of U1 snRNA, indicating that each nucleotide position is critical for the chromatin association of NXF1-enChr RNA. In another example, the ‘G’ to ‘C’ mutation at the 73G position outside the 7-nt U1 recognition motif caused a 3.2-fold increase of the cyto/chr ratio (Figure 2A). Structural analysis of the 73G>C mutation suggested shifted base-pairing in the stem and an abnormal loop sequence, which contains only 2 nucleotides of the 7-nt sequence (Figure 2D). Sequestration of this 7-nt U1 recognition site into a stem region in the 73G>C mutant likely led to impaired chromatin association of *NXF1-enChr* RNA (Figures 2A and 2D).

In comparison, mutations at the 42G and 51G positions of NXF1-enChr led to subtle decreases in the cyto/chr ratio (0.90 and 0.76, respectively), indicating enhanced chromatin association. Interestingly, compared to ‘42G’ in the wild-type sequence, the ‘42G>A’ mutant appears to base pair perfectly with the fourth ‘U’ nucleotide of the U1 targeting sequence (Figures 2A and 2C). In the 51G>C mutant, shifted sequence pairing in the stem region may lead to nearly full exposure of the 9- nt U1 recognition motif (nucleotides 38 to 45), thereby enabling better U1 binding. Additionally, a weak U1 recognition site at positions 53 to 59, which is partially exposed in the two middle bulges in the wild-type structure (marked by a thin blue line in Figures 2B and S3E, Figure S3G), now appears to be better exposed in the 51G>C mutant (Figure 2E). Mutations or deletions within this weak U1 site also slightly decrease the chromatin association of NXF1-enChr RNA, with ~1.5-2-fold increases of the cyto/chr ratio (Figure 2A). Thus, perfect base-pairing or better exposure of the U1 recognition sites in the 42G>A and 51G>C mutants may promote efficient binding of U1 snRNA, consequently leading to enhanced chromatin localization of NXF1-enChr (Figures 2A and 2E).

In summary, mutREL-seq revealed a novel U1 recognition motif required for chromatin retention of NXF1-enChr RNA. Increased or disrupted base-pairing with U1 snRNA correlates with enhanced or decreased chromatin association of *NXF1-enChr* RNA, respectively (Figure 2A). The opposing effects of the 51G>C and 73G>C mutations suggested that the sequences surrounding the U1 recognition motifs also contribute to efficient U1 binding and RNA chromatin localization, probably by making the U1 sites available for U1 snRNP recognition and binding.

### U1 recognition motif and binding are enriched in enChrs

Next we asked whether the U1 motif and its binding are enriched in the enChrs we identified. We performed bioinformatics prediction of U1 recognition sites with strong or medium strength across human and mouse transcriptomes. We also analyzed the RNA interactome of U1 snRNA that was previously revealed by U1 RAP-RNA-seq (RNA affinity purification followed by RNA sequencing) in mESCs (Engreitz et al., 2014). The majority of enChrs overlap well with predicted U1 recognition sites and/or U1 RAP-RNA signals except *Xist* enChrs (Figures 1C, 1D, S2, and Table S2). For example, multiple strong U1 recognition sites are clustered in the 162-nt NXF1-enChr and nearby sequences. U1 snRNP also binds strongly (~70-fold maximal enrichment versus the input) to this region with peak signals that are centered at the 7-nt U1 motif revealed by mutREL-seq (Figures 1D and 2A), supporting direct interactions of *NXF1-enChr* RNA and U1 snRNP *in vivo.*

For mouse *Malat1,* multiple enChrs were found within mouse repeats E and M which harbor multiple U1 recognition sites and exhibit high-level binding of U1 snRNP (~35- or 42-fold maximal enrichments versus the input) (Figure 1C and Table S2). In fact, *U1:Malat1* interactions have reportedly been conserved between human and mouse (Lu et al., 2016). Thus, U1 targeting may constitute a conserved mechanism that controls *Malat1* nuclear and chromatin retention across mammalian species. Compared to simulated controls, enChrs exhibit 2.6-4-fold higher overlaps with predicted U1 recognition sites (Figure 2F and Table S2). In comparison, for cytoplasm-localized protein-coding transcripts, U1 signals are mainly confined in the intronic regions and appear to be depleted in exons (Figures S1F-S1H, S3H, and S3I), which is consistent with a previous study (Engreitz et al., 2014).

### A U1 recognition motif retains *GFP* reporter RNA on chromatin

To test whether the U1-containing enChr (U1) affects the subcellular localization of a *GFP* reporter RNA, we inserted the wild-type and mutant NXF1-enChr between a *GFP* gene and a polyadenylation signal (PAS) from the bovine growth hormone gene (Figure 3A). Mutation (mutU1) of the 7-nt of the core U1 recognition site did not change the overall structure of NXF1-enChr (Figure S4A). We also confirmed that the insertion of *NXF1-enChr* (U1) did not elicit splicing (Figure S4B). Interestingly, the NXF1-enChr (U1) strongly suppressed GFP fluorescence signals, and retained ~40% of *GFP* transcripts in the chromatin fraction at a level comparable to that of *Malat1* (Figures 3B and 3C). In contrast, mutation of the U1 site completely abolished the chromatin association of *GFP* RNA and rescued GFP protein expression (Figures 3B and 3D). Intriguingly, we found that insertion of the wild-type but not mutant enChr (U1) significantly reduced the level of *GFP* transcripts to ~20% of that of cells expressing *GFP* alone (Figure 3D).

**Figure 3.**
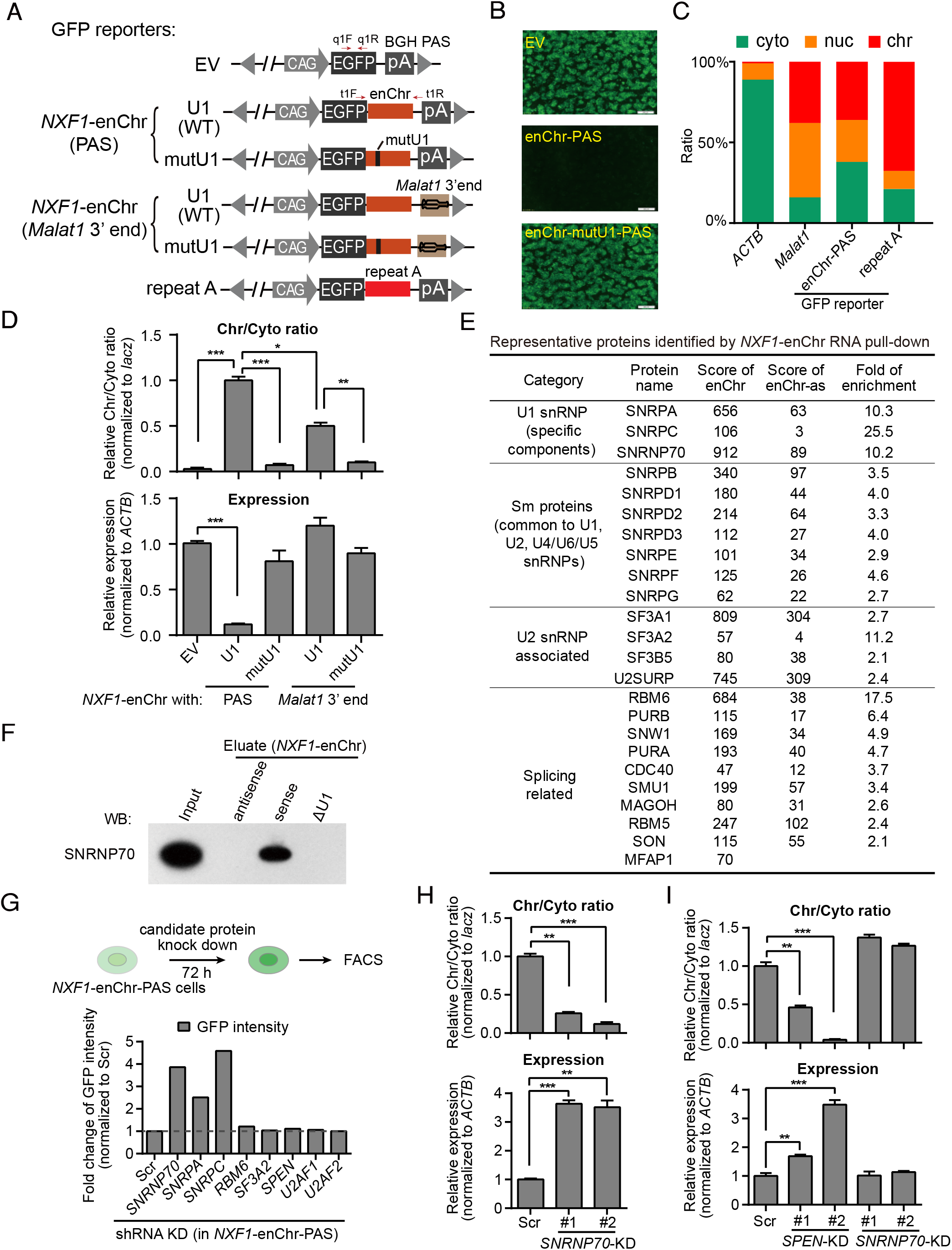
U1 snRNP directly regulates RNA chromatin retention. (A) The schematic diagrams show the information of GFP reporters used. The two pairs of red arrows show the relative positions of the primers used to analyze potential splicing of the insert (t1F/t1R, related to Figure S4B) and RT-qPCR analysis (q1F/q1R, related to Figures 3C, 3D, 3H, and 3I). EV, empty vector; BGH PAS, bovine growth hormone polyadenylation signal; *Malat1* 3’ end, 3’ termination sequence of *Malat1,* from the putative nuclear retention element (ENE) to the end of *Malat1;* WT, wild-type; mutU1, mutated U1 recognition site in NXF1-enChr, show in vertical black line. (B) GFP fluorescence imaging of ES cells expressed the EV, NXF1-enChr-PAS, or NXF1-enChr-mutU1-PAS constructs as shown in Figure 3A. (C) RT-qPCR analysis of the subcellular localization of GFP-NXF1-enChr fused RNA and control genes *ACTB* and *Malat1.* (D) RT-qPCR analysis of the chr/cyto ratio (top panel) and relative expression (bottom panel) of GFP-NXF1-enChr fused RNA identified from subcellular fractionation in cells expressed different constructs as shown in Figure 3A. (E) Representative proteins identified by the *NXF1-enChr* RNA pull-down assay. The mass spectrometry score of the proteins identified by the *NXF1-enChr* and NXF1-enChr-as RNA pull-down assays are shown, together with their fold enrichment in the NXF1-enChr sample relative to the NXF1-enChr-as sample. (F) Western blot to confirm the specific interaction between SNRNP70 and NXF1-enChr. Antisense sequence and sequence with the deletion of the strong U1 recognition site (ΔU1) were used as controls. (G) Schematic diagram shows the procedure of the mini-screen (top panel) and fold change of GFP intensity after different proteins depletion in *NXF1*-enChr-PAS GFP reporter cells (bottom panel). (H) RT-qPCR analysis of the chr/cyto ratio (top panel) and relative expression (bottom panel) of GFP-*NXF1* -enChr fused RNA after *SNRNP70* knock down. (I) RT-qPCR analysis of the chr/cyto ratio (top panel) and relative expression (bottom panel) of GFP-repeat A fused RNA after *SPEN* or *SNRNP70* knock down. The relative chr/cyto ratio was normalized to an in vitro transcribed *lacZ* RNA, which was used as a spike-in during subcellular fractionation. the relative expression level change was normalized to *ACTB.* Data are shown as mean ± sem. including two biological replicates and two technical replicates. * *p* < 0.05, ** *p* < 0.01, *** *p* < 0.001.

It was reported that U1 inhibits the PAS and polyadenylation to promote pre-mRNA degradation (Fortes et al., 2003; Goraczniak et al., 2009; Gunderson et al., 1998). To rule out an indirect effect of RNA degradation on its chromatin localization, we replaced the PAS with the 3’ termination sequence of *Malat1* (Figure 3A). The 3’ end of *Malat1* possesses a triple-helix structure, which resembles the viral expression and nuclear retention element (ENE) and stabilizes the *Malat1* transcripts (Brown et al., 2012; Wilusz et al., 2012). Replacement of the PAS with the *Malat1* 3’ termination sequence abolished the inhibitory effect of the enChr (U1)-PAS axis and rescued the expression of *GFP* RNA. Consistently, the wild-type but not mutU1 enChr significantly promoted the chromatin association of *GFP* transcripts despite comparable RNA levels (Figure 3D). These results indicate that the U1 recognition motif directly contributes to the chromatin association of *GFP* reporter RNA regardless of RNA stability.

Notably, GFP-enChr (U1) RNA was expressed ~5-fold higher in the 3’ *Malat1* reporter compared to its expression in the PAS reporter. However, the chr/cyto ratio of GFP-enChr (U1) RNA was ~1-fold lower in the 3’ *Malat1* reporter (Figure 3D). This result supports the notion that RNA turnover and instability may promote the binding of RNA to chromatin. Thus, the U1-PAS axis might also contribute to the enriched pattern of chromatin associations for unstable RNA transcripts by promoting RNA degradation.

### U1 snRNP is required for the chromatin retention of *GFP* reporter RNA

To reveal proteins that are involved in RNA-chromatin retention, we used NXF1-enChr (U1) RNA as the bait to pull down its interacting proteins for mass spectrometry analysis (Figures S4C and S4D). Among 79 proteins that specifically interact with the NXF1-enChr RNA (mass-spec score > 12 and enrichment of sense versus antisense > 2-fold), 41 proteins are known to function in splicing and 14 are core components of the spliceosome (Figure 3E and Table S3). This list includes the U1 snRNP-specific components SNRPA, SNRPC and SNRNP70 that are involved in pre-catalytic spliceosome assembly, and a number of general Sm proteins, such as SNRPDs (1, 2 and 3) and SNRPs (B, E, F and G) that mediate the assembly of multiple U1, U2 snRNPs, and U4/U6 and U5 tri-snRNPs, and the splicing cofactors RBM6, SON, and CDC40 (Ahn et al., 2011; Bechara et al., 2013; Ben Yehuda et al., 1998; Will and Luhrmann, 2011). We noted that components specific for catalytic snRNPs were not detected by NXF1-enChr RNA pull-down (Figure 3E and Table S3). Western blotting analysis confirmed that the enChr RNA, but not the antisense or a mutant RNA with the 9-nt U1 site deleted (enChr-ΔU1), captured SNRNP70 (Figure 3F). Thus, U1 snRNP binds to the *NXF1-* enChr via the U1 recognition site *in vitro.*

We then selected a subset of enChr-interacting proteins and analyzed the effects of their knockdown on the expression and chromatin association of *GFP* RNA by using the enChr (U1)-PAS reporter (Figures 3E, 3G, and S4E). To assess the specificity of U1-mediated chromatin association, we constructed another *GFP*-repeat A reporter by inserting the repeat A sequence of *Xist* between a *GFP* and PAS (Figure 3A). We selected the repeat A because it was identified as a chromatin enriched fragment by REL-seq but does not harbor a U1 recognition site. The repeat A is known to recruit the repressor protein SPEN (also known as SHARP) which is essential for XCI (Figures 3C and S2A) (Chu et al., 2015; McHugh et al., 2015; Monfort et al., 2015). Using the GFP-enChr (U1) reporter, we found that depletion of the three core components of U1 snRNP *(SNRNP70, SNRPA, SNRPC),* but not *SPEN* and splicing regulators *(RBM6, SF3A2, U2AF1/2),* increased GFP fluorescence signals by 2.5-4.5-fold (Figure 3G). Knockdown of *SNRNP70* dramatically increased the expression but decreased the chr/cyto ratio of GFP-enChr (U1) RNA, a pattern mimicking the effect of mutating the U1 site of NXF1-enChr (Figures 3H and S4F). However, *SNRNP70* knockdown failed to affect the expression and chromatin location of GFP-repeat A RNA in contrast to depletion of *SPEN* (Figure 3I). These results indicated that U1 snRNP specifically promotes the chromatin association of a *GFP* reporter RNA that harbors the U1 recognition site.

### U1 motifs and snRNA binding are globally enriched in chromatin-bound lncRNAs

Having confirmed a direct involvement of the U1 recognition motif and U1 snRNP in regulating the chromatin localization of a reporter RNA, we then sought to study the biological significance of U1 targeting on RNA-chromatin association in the cell. We first examined the global correlation of U1 recognition and binding with RNA chromatin localization. Higher densities of U1 recognition motifs (strong or medium) across exons were observed in all lncRNA transcripts compared to protein-coding mRNAs regardless of their expression in mouse and human (Figures S5A and S5B). This pattern of U1 site distribution became more obvious when comparing the top 20% chromatin-enriched lncRNAs with the top 20% chromatin-depleted mRNAs (Figures 4A and 4B).

**Figure 4.**
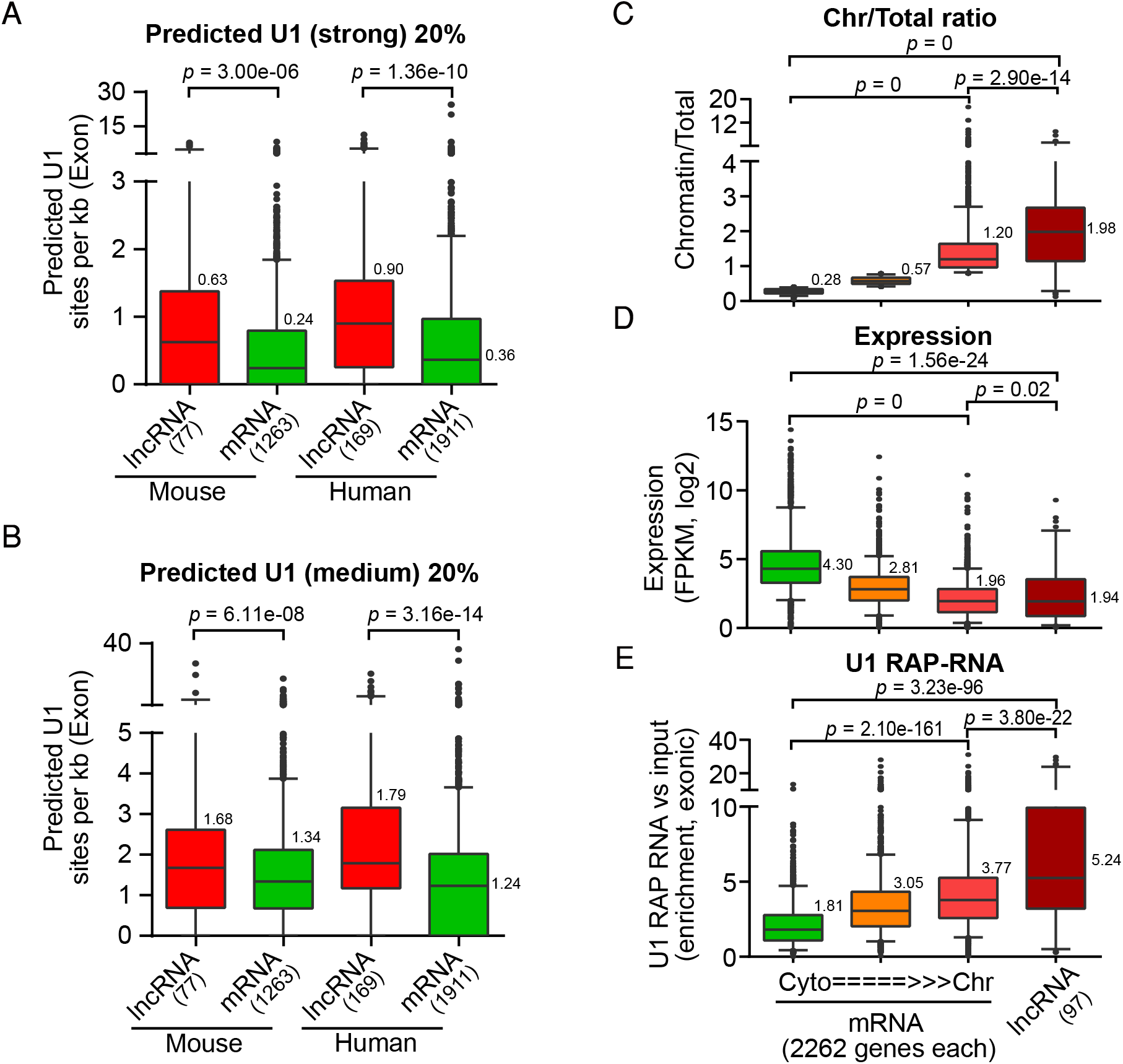
U1 motifs and U1 snRNA binding are globally enriched in chromatin-bound lncRNAs. (A, B) Comparison of the density of predicted strong (A) or medium (B) U1 recognition site in exons between the top 20% of chromatin-enriched lncRNAs and the top 20% of chromatin-depleted coding genes in human and mouse. (C-E) Comparison of chromatin enrichment (C), expression (D), and fold enrichment of U1 RAP-RNA signal (E) among coding genes with different chromatin association tendency and lncRNAs. T-tests were used to compare the data for two indicated samples. The median value for each sample is labeled beside the box-plot. As the RNA-seq of subcellular fractionation is un-stranded, we discarded lncRNA transcripts that overlapped with protein coding genes for the analysis.

Next we took 97 lncRNAs and 6786 mRNA transcripts that show no overlap in their genomic sequences and can be detected both in mESCs (FPKM > 1) and by U1 RAP-RNA-seq (> 10 exonic reads in both U1 RAP-RNA and the input). We further divided mRNA transcripts into three groups based on their subcellular localization. The group of 97 lncRNAs shows a significantly higher median value of the chromatin versus total cellular RNA ratio compared to the group of the most chromatin-enriched mRNAs (1.98 versus 1.20, *p* = 2.9e-14) (Figure 4C). Interestingly, levels of U1 binding in exons gradually increase from the most cytoplasm-enriched RNAs to chromatin-enriched mRNA and then to lncRNAs (median enrichments from 1.81 to 3.77 and to 5.24, respectively), whereas their expression decreases along with increased chromatin association (Figures 4D and 4E). This result indicates a general positive correlation of RNA-chromatin association with U1 binding but an inverse correlation with gene expression.

In contrast to exons, introns show comparable, high-level U1 binding (median enrichment 6.12 - 8.58) regardless of whether they are located in lncRNAs and mRNAs (Figure S5C). This observation is consistent with the fact that most intronic RNAs are quickly degraded upon splicing and rarely leave the nucleus. Together, the globally enriched distribution of U1 recognition sites and the genome-wide high-level binding of U1 snRNA on chromatin-associated transcripts suggest a broad role of U1 snRNA in the regulation of RNA chromatin localization.

### U1 inhibition downregulates global binding of lncRNAs on chromatin

To investigate the direct role of U1 snRNP in regulating lncRNA-chromatin association, we then sought to inhibit U1 by antisense morpholino oligonucleotide (AMO) in mESCs. To explore the possible involvement of splicing in RNA-chromatin association, we also sought to inhibit U2 snRNA or both U1 and U2 (U1/2). Considering the broad involvement of U1 and U2 snRNAs in cellular functions (Clark et al., 2012), we performed a short-term treatment with U1 and/or U2 AMOs in order to assess the immediate effects on RNA subcellular localization. Short-term treatments appeared to be suitable for studying lncRNA and ncRNA transcripts with relatively short half-lives (see below and discussion).

We first examined several chromatin-bound lncRNAs that exhibit high levels of U1 RAP-RNA signals on their transcripts, including *Malat1, Neat1, NR_028425, Pvt1, Kcnq1ot1,* and *Tsix* (Figures 5A and S6A). Inhibition of U1 alone or both U1/2 significantly reduced the chr/cyto ratios of these lncRNAs, while inhibition of U2 alone had a much subtler effect. Notably, inhibition of both U1/2 did not result in a stronger effect than inhibition of U1 alone, indicating a major role for U1 snRNA in regulating the chromatin association of these lncRNAs. Next, we tried to knock down *SNRNP70* and observed efficient depletion after 3 days of treatment with shRNAs (Figure S4F). Depletion of *SNRNP70* downregulated RNA-chromatin associations, except for *Neat1* which might be indirectly upregulated due to the prolonged treatment (Figures 5B and S6B). Consistent with the fact that *Malat1* is more stable than other lncRNAs (Clark et al., 2012), inhibition of *SNRNP70* caused a more dramatic reduction of *Malat1* on chromatin than short-term inhibition of U1/2 (Figures 5B and S6B).

**Figure 5.**
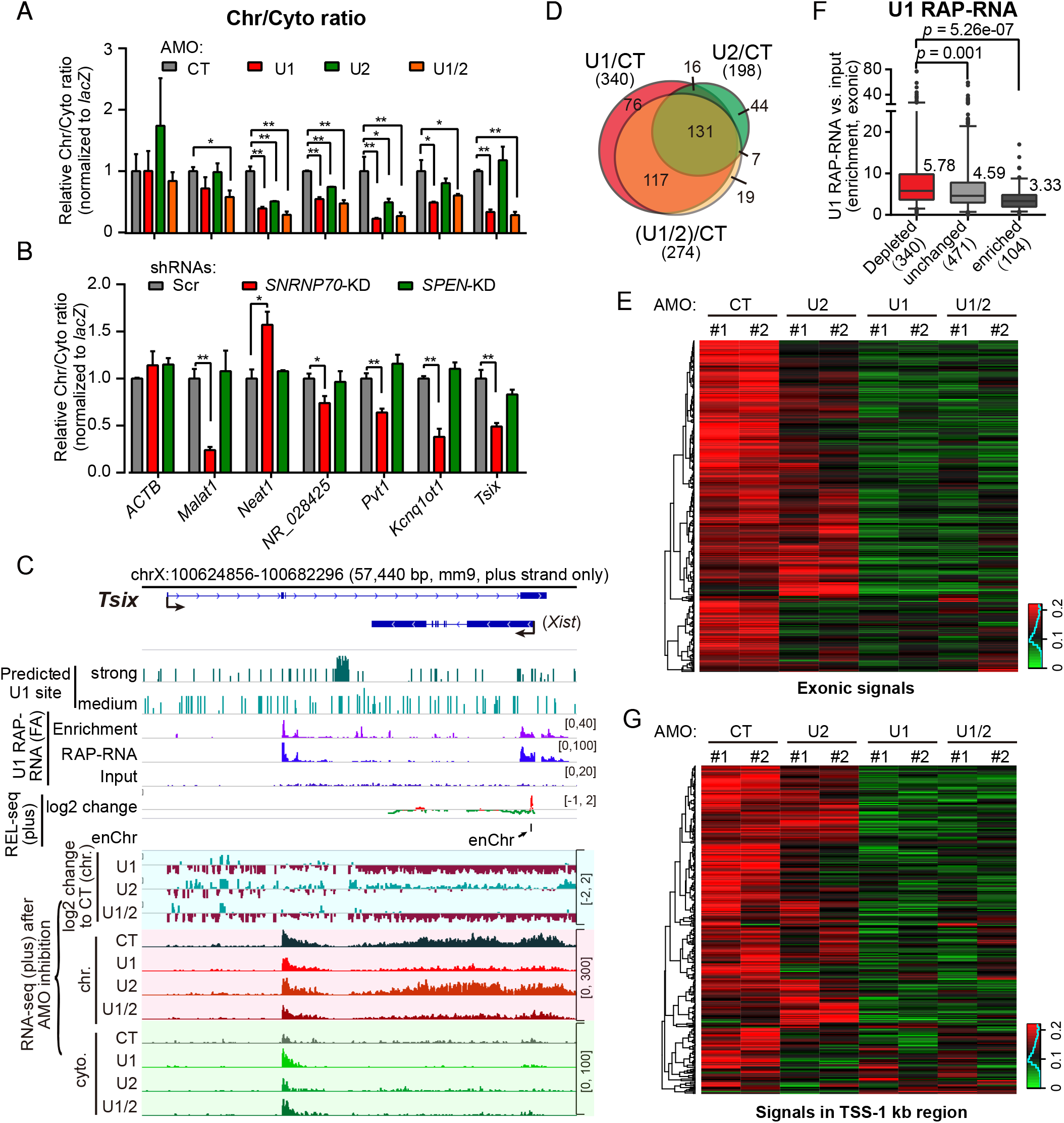
U1 snRNP regulates the chromatin retention of lncRNAs. (A, B) RT-qPCR analyses of the relative chr/cyto ratios of candidate lncRNAs after short-term treatment by control AMO (CT), U1, U2, or both U1/2 AMOs (A), or after scramble (Scr), *SNRNP70*, or *SPEN* shRNA knock down (B). Data are shown as mean ± sem. including two biological replicates and two technical replicates. * *p* < 0.05, *\*\*p* < 0.01. (C) Sequencing tracks of chromatin and cytoplasmic RNA-seq signals at *Tsix* locus after U1, U2, U1/2 inhibition. The enChr identified by REL-seq is highlighted. Only the tracks of plus strand are shown. (D) Venn diagram of lncRNAs with decreased chromatin association upon U1, U2, or U1/2 inhibition ((U1 or U2 or U1/2) versus CT < 1, *p* < 0.05 of two biological replicates, t-test). (E) Heatmap of chromatin RNA-seq signals of 340 lncRNAs with decreased chromatin association upon U1 inhibition. (F) U1 RAP-RNA signals in the sets of lncRNAs that show depleted, enriched or unchanged levels on chromatin in Figure S6H. The median values were shown. (G) Heatmap of chromatin RNA-seq signals in the TSS-1 kb region for 307 lncRNAs that show detectable RNA reads (RPM in TSS-1 kb > 0.5) out of 340 lncRNAs with decreased chromatin association shown in panel (E).

To assess the genome-wide effects of U1 and/or U2 inhibition on RNA subcellular localization, we performed strand-specific RNA-seq analysis of total RNAs in chromatin and cytoplasmic fractions. Increased accumulation of intron-retaining mRNA transcripts in the cytoplasmic fraction indicated the effectiveness of splicing inhibition by 2-hour treatments with U1 and/or U2 AMOs (Figures S6C and S6D). Consistent with the analysis of individual lncRNAs, inhibition of U1 or both U1/2 had more dramatic effects than inhibition of U2 (e.g., *Tsix* and *Pvt1,* Figures 5C and S6E). Compared to the control AMO (CT; *p* < 0.05, two biological replicates), U1 AMO downregulated the binding of 340 lncRNAs on chromatin, while inhibition of both U1/2 affected 274 lncRNAs. There was extensive overlap of the lncRNAs between these two treatments. In comparison, U2 AMO alone only affected the binding of 198 lncRNAs on chromatin, the majority (74%) of which showed decreased chromatin binding upon U1 inhibition (e.g., *Neat1* and *NR_028425;* Figures 5D, S6F and S6G). Heatmap analysis further revealed that the degrees of downregulation upon U2 inhibition were much more subtle than those caused by inhibition of U1 or both U1/2 (Figure 5E). These results illustrated that U1 snRNA plays a more dominant role than U2 in tethering lncRNAs on chromatin.

On the other hand, the fact that U2 inhibition also altered the localization of a subset of chromatin-bound lncRNAs, albeit to a lesser degree than U1 inhibition, suggested that splicing might be involved in the regulation of lncRNA-chromatin association.

Upon U1 inhibition, more lncRNAs showed decreased than increased chromatin associations (340 versus 104) (Figure S6H). In addition, the set of 340 lncRNAs with decreased chromatin associations exhibited higher U1 RAP-RNA signals compared to lncRNAs with no change (471) or with upregulated chromatin occupancies (104) (median-level enrichment: 5.78 versus 4.59 and 3.33, respectively) (Figure 5F). Next, to rule out the possibility that lncRNA expression might affect its chromatin localization, we compared the chr/cyto ratio of U1 and/or U2 inhibition versus the control. Consistent with the analysis of absolute RNA signals on chromatin, we observed a similar global downregulation of lncRNA chromatin associations (Figure S6I). Particularly for the set of 340 lncRNAs with decreased chromatin associations upon U1 inhibition, the majority (229) had a reduced chr/cyto ratio [U1 versus CT (chr/cyto) < 0.8]. This result indicated that the observed decreases of chromatin signals were not simply due to reduced levels of total transcripts (Figure S6I).

It was reported that U1 knockdown terminated most nascent transcripts at ~1 kb from the start of transcription, whereas splicing inhibition by U2 AMO or spliceostatin A (SSA), which specifically inactivates the U2 snRNP component SF3B, had no effect despite the fact that inhibition of all these factors efficiently blocked splicing (Berg et al., 2012; Kaida et al., 2010). To minimize the possible influence of pre-mature termination induced by U1 inhibition on the analysis of chromatin-bound lncRNAs, we analyzed chromatin RNA-seq signals in the 5’ end of the transcript from the TSS to the 1-kb downstream region, in which RNA levels are less likely to be affected by U1 inhibition (e.g., *DTNBP1,* Figure S6J). Among 307 lncRNAs that were downregulated on chromatin and showed detectable RNA signals in their 5’ region (RPM > 0.5) upon U1 inhibition, the majority (270 with U1/CT < 0.8; or 196 with U1/CT < 1 and *p* < 0.05) exhibited decreases of chromatin-associated RNA reads in their 5’ regions (Figures 5G and S6K). Thus, the global downregulation of lncRNA-chromatin association upon U1 or both U1/2 inhibition probably did not result from the effect of U1 telescripting. Together, these results demonstrated that U1 snRNA, and to a less degree U2, directly regulates the chromatin association of lncRNAs, independent of transcription termination and total RNA levels.

### Inhibition of U1 and U2 downregulates chromatin associations of PROMPTs and eRNAs

Regulatory DNA elements produce many chromatin-associated ncRNA transcripts, including eRNAs and PROMPTs. Because of their unstable nature and low abundance in cells, few RNA reads of U1 RAP-RNA sequencing fall into these transcripts. U1 RAP-DNA sequencing has revealed that U1 binds to the TSSs of active genes independent of transcription elongation (Engreitz et al., 2014). Intriguingly, in addition to the reported binding at the TSSs, we found that U1 snRNA also binds to enhancers and super-enhancers (Figure S7A). The enrichment of U1 in the genomic neighborhood of PROMPTs and eRNAs supports a functional interaction between U1 and these ncRNAs. Indeed, visual inspection of individual loci showed altered chromatin associations of eRNAs and PROMPTs upon inhibition of U1 and/or U2. For example, in an upstream ~10-kb promoter region covering a super-enhancer of *PHC1,* antisense transcripts representing *PHC1* eRNA or PROMPT were robustly detected in mESCs. Inhibition of U1 or both U1/2 led to decreased chromatin signals and increased cytosolic signals of *PHC1* eRNA/PROMPT. In contrast, signals of chromatin-bound *PHC1* pre-mRNA were increased, likely reflecting defective splicing and abnormal intron retention upon inhibition of U1 or U1/2 (Figure 6A). In another example, an enhancer located ~23 kb upstream of the lncRNA *Haunt* produces sense and antisense eRNA transcripts which were detected in the chromatin but not the cytosolic fraction in control mESCs. Inhibition of U1 or both U1/2 dramatically increased the cytosolic signals of the sense and antisense eRNA transcripts, implying increased nuclear export (Figure 6B).

**Figure 6.**
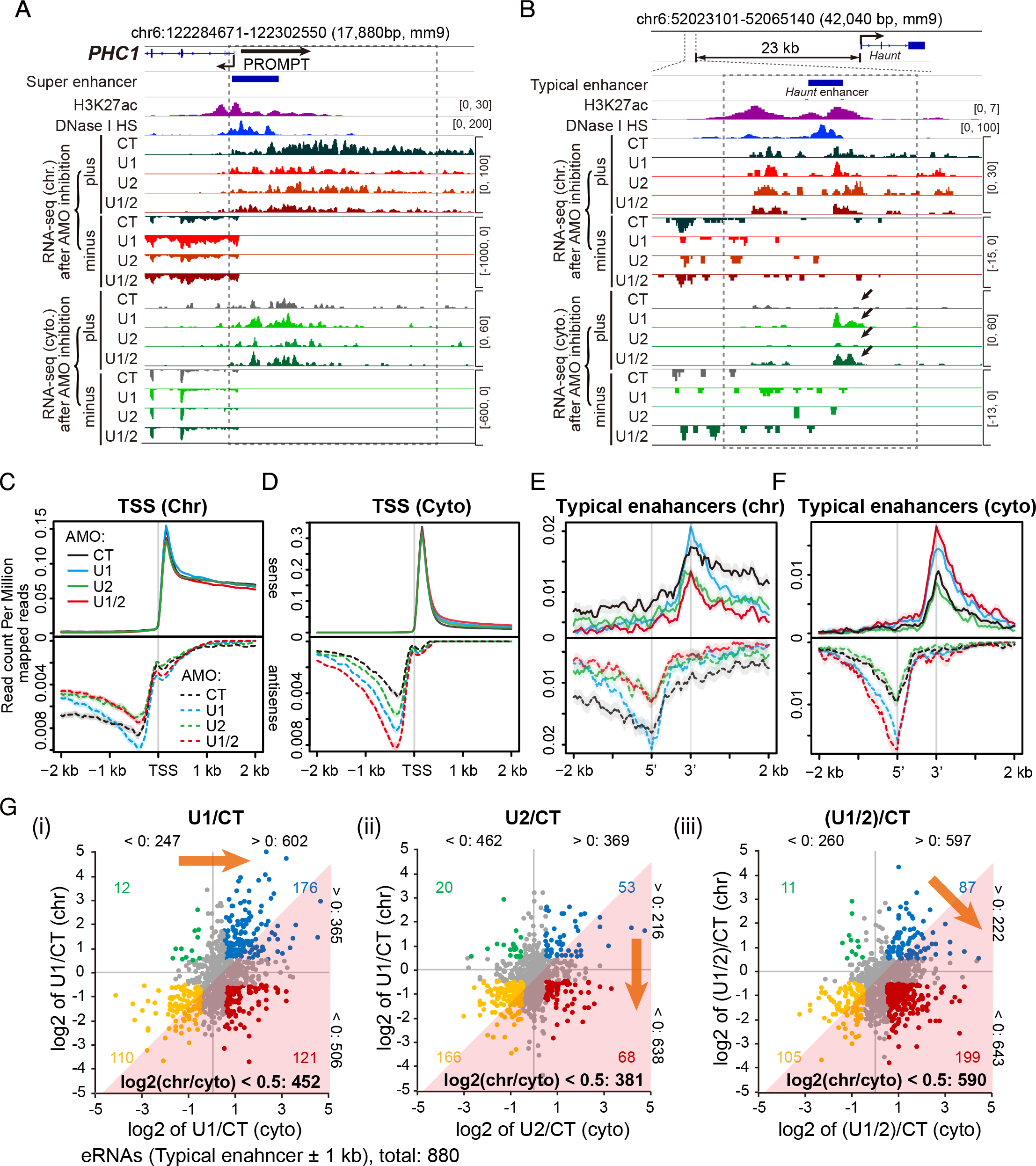
Inhibition of U1 and U2 downregulates chromatin associations of PROMPTs and eRNAs. (A, B) Sequencing tracks showing chromatin and cytoplasmic RNA-seq signals of a PROMPT/eRNA in the *PHC1* locus (A) and eRNAs in the *Haunt* enhancer region (B). Dashed boxes highlight the PROMPT (A) or the eRNA (B) region. Black arrows in (B) highlight the increased cytoplasmic signal upon U1 or U1/2 inhibition. (C, D) Metaplots of chromatin (C) or cytoplasmic (D) RNA-seq reads in CT (black line), U1 (blue), U2 (green), and U1/2 (red) AMO-treated samples in a ±2 kb window flanking the TSS of all *Ensembl* genes. Dashed lines represent the plots of antisense reads from the respective samples, and solid lines represent sense signals. Shaded regions represent the standard error of the mean. (E, F) Same as (C, D) for 880 typical enhancers that show detectable eRNA expression and do not overlap with any genes within 2 kb. (G) Scatter plots show the fold change (log2) of RNA-seq signals in the chromatin or cytoplasmic fraction of 880 typical enhancers (the same set as shown in Figures 6E and 6F) upon U1 (i), U2 (ii), or U1/2 (iii) inhibition. Each dot represents the relative change of an eRNA in the chromatin (y-axis) or cytosolic (x-axis) fraction compared to the control (CT). RNA reads in a region from 1 kb upstream to 1 kb downstream of each enhancer were taken into account. Regions within the pink triangles comprise eRNAs with decreased chr/cyto ratios, as judged by comparing U1 and/or U2 AMO-treated samples to the control [log2 of ((U1 and/or U2)/CT (chr/cyto)) < 0.5]. Orange arrows represent the overall tendency of changes in chromatin and cytoplasmic locations. eRNAs with log2 of fold-change > 0.5 or < −0.5 in cytoplasmic or chromatin fractions are highlighted with colors, and the corresponding numbers of changed eRNAs were shown in each quadrant. The number of eRNAs that show increased (log2 of fold-change > 0) or decreased (log2 of fold-change < 0) changes in chromatin fraction or cytoplasmic fraction are labeled in black (eRNAs with log2 of change to CT = 0 were not taken into account).

To assess genome-wide changes in the subcellular localization of ncRNAs upon AMO treatments, we performed metagene analyses of RNA-seq reads near the TSSs of all *Ensembl* genes for PROMPTs, and at a set of 880 typical enhancers that show detectable RNA signals and do not overlap a gene within a 2-kb range for eRNAs. Compared to inhibition of either U1 or U2 alone, inhibition of both U1/2 caused more dramatic decreases in the chromatin signals and increases in the cytosolic signals of PROMPTs (antisense) and eRNAs (both sense and antisense). In comparison, U1 inhibition significantly increased the cytosolic levels of PROMPTs and eRNAs, and downregulated their levels in the chromatin beyond 1-kb upstream of the TSS or surrounding the 5’ and 3’ boundary of enhancers, but had a minor effect on chromatin-RNA levels within 1-kb regions (Figures 6C-F). U2 inhibition downregulated the chromatin signals of both classes of ncRNA, and slightly increased cytosolic signals for PROMPTs, but had no effect on cytosolic eRNAs. Similar patterns were observed for eRNAs produced at 100 mESC-specific super-enhancers that do not overlap with any genes (Figures S7B and S7C).

We then calculated changes in the chr/cyto ratio in U1 and/or U2 AMO-treated samples to that of the control for eRNAs [log2 of (U1 and/or U2)/CT (chr/cyto) < 0.5; regions shaded pink in Figure 6G]. Remarkably, out of 880 eRNAs analyzed, 590 showed a decreased ratio of chromatin versus cytosol fractions upon inhibition of both U1/2, while 452 and 381 eRNAs showed decreases upon inhibition of either U1 or U2, respectively (Figure 6G). Inhibition of U1 or U2 alone led to a shift in the distribution of eRNA localization towards the bottom of the plot [log2 of (U1 or U2)/CT (chr) < 0] (Panels i and ii of Figure 6G). However, a more notable change upon U1 inhibition is the increase of cytosolic RNA signals as shown by a shifted distribution of eRNA localization towards the right of the plot [log2 of U1/CT (cyto) > 0] (Panel i of Figure 6G). Inhibition of both U1/2 led to dramatic downregulation of the absolute and relative chromatin signals of eRNAs, resulting in a shifted distribution towards the bottom right of the plot [log2 of (U1/2)/CT (cyto) > 0 and log2 of (U1/2)/CT (chr) < 0] (Panel iii of Figure 6G).

Thus, both U1 and U2 snRNAs directly promote the chromatin association of a subset of PROMPTs and eRNAs. In addition, U1 snRNA appears to inhibit the nuclear export and cytosolic expression of PROMPTs and eRNAs. Together, U1 and U2 snRNAs act synergistically to tether PROMPTs and eRNAs on chromatin and to prevent their export to the cytosol.

## Discussion

Thousands of functionally uncharacterized noncoding transcripts exist, and they are commonly found in chromatin fraction. We hypothesized that cis-elements embedded in the DNA or RNA transcript may determine the binding of specific *trans* factor, thus contributing to RNA subcellular localization. To systematically carry out high-resolution screens for cis-regulatory RNA sequences that contribute to RNA chromatin localization, we developed a novel technique named REL-seq. As a proof of principle, we analyzed 9 lncRNA and mRNA transcripts and identified 26 enChrs that are 50-500-nt in length and direct strong chromatin enrichment of the host RNA, demonstrating the effectiveness of the REL-seq strategy. During the preparation of this manuscript, two papers were published which used a similar screening strategy and identified Alu repeat-like sequences and a number of RNA nuclear enrichment sequences (Lubelsky and Ulitsky, 2018; Shukla et al., 2018). In this study, to identify key residues that directly regulate RNA chromatin localization, we further developed mutREL-seq by combining REL-seq with random mutagenesis. Importantly, mutREL-seq analysis of an enChr in the intron-retaining isoform of *NXF1* revealed a 7-nt U1 recognition motif critical for its retention on chromatin. Reporter assays demonstrated a key role for the U1 motif and U1 snRNP in regulating the expression and chromatin association of a *GFP* reporter RNA. Interestingly, lncRNAs and chromatin-enriched mRNAs show genome-wide enrichments of U1 recognition motifs and high-level direct binding of U1 snRNA on their RNA transcripts in both human and mouse. Remarkably, short-term inhibition of U1 led to globally downregulated binding of lncRNAs on chromatin, while inhibition of both U1 and U2 caused more dramatic defects in the chromatin localization of PROMPTs and eRNAs compared to inhibition of either U1 or U2. In summary, we propose that U1 snRNP, and perhaps the splicing machinery, act widely to promote the chromatin association of noncoding transcripts.

Recognition of the 5’ splice site by U1 snRNP has been implicated in regulating nuclear retention of intron-retaining mRNA transcripts. Splicing inhibition by SSA causes widespread intron retention and pre-mRNA accumulation in nuclear speckles (Kaida et al., 2007; Martins et al., 2011; Yoshimoto et al., 2017). Intriguingly, a small number of short pre-mRNAs, which fail to be recognized by U1 snRNP due to weak 5’ splice sites, leak into the cytoplasm, indicating a role of U1 recognition in retaining pre-mRNAs in the nucleus (Yoshimoto et al., 2017). In addition, the U1 snRNP component SNRNP70 was reported to be required for the nuclear retention of *p27* pre-mRNAs (Takemura et al., 2011). A study utilizing the *fushi tarazu (ftz)* mini gene reported that the presence of a 5’ splice site motif inhibited *ftz* mRNA nuclear export, 3’ cleavage and polyadenylation, yet promoted *ftz* mRNA degradation (Lee et al., 2015). In addition to the canonical U1 motif, the 5’ splice site recognized by U11 snRNP, a minor spliceosome recognition complex equivalent to the major U1 snRNP, was reported to promote the nuclear retention of *U11/U12-65K* mRNA (Patel and Steitz, 2003; Verbeeren et al., 2017). Here using a distinct, unbiased screening approach, we discovered that the U1 recognition motif regulates the chromatin localization of noncoding RNAs. Together with studies on a handful of mRNA transcripts, our findings indicate that U1 recognition may be universally employed by many RNA transcripts to regulate their chromatin association and expression.

It should be noted that not all predicted U1 sites were identified as chromatin-enriched signals by REL-seq, and not all enChrs contain U1 recognition sites. First, not all predicted U1 recognition sites are bound by U1 snRNP as shown by U1 RAP-RNA-seq (Figures 1C, 1D and S2). Opposing effects observed in the NXF1-enChr 51G>C and 73G>C mutants indicate that the peripheral sequences are required to facilitate U1:RNA base pairing by stabilizing a local RNA structure to present the U1 recognition site for base pairing (Figure 2). In addition, U1 snRNP and splicing-related RNA-binding proteins including U2AFs and SR proteins bind to broad regions of enChrs and nearby sequences beyond the core 7-nt U1 motif (Figures 2A, S4E). The likelihood of U1 binding may be enhanced by synergistic interactions of multiple proteins on the peripheral sequences, therefore reinforcing the chromatin retention effect of U1 snRNP and associated factors. Second, we believe that U1 snRNP targeting is prevalent, but not the only mechanism for RNA nuclear retention. Repetitive sequences have been implicated in regulating chromatin binding of the lncRNAs *Xist* and *Firre*. We identified all four repeat regions that were previously reported as important for *Xist* chromatin association and XCI (Chen et al., 2016; Lv et al., 2016; Wutz et al., 2002). Fusion of repeat A to a *GFP* RNA led to its chromatin retention in a manner that depended on *SPEN* rather than *SNRNP70* (Figures 3A, 3C, and 3I). In another example, *Firre* transcripts contain 8-16 copies of a unique, conserved 156-nt repeating RNA domain (RRD) that is bound by HNRNPU (Hacisuleyman et al., 2016). Both the RRD and HNRNPU are required for the proper focal and nuclear localization of *Firre.*

Most of the enChrs do not harbor splice acceptor sites, and no splicing was detected in the *NXF1*-enChr region (Figure S4B). These results indicate that the enChrs consist of cryptic 5’ splice sites, which may serve a non-splicing purpose to recruit U1 snRNP to mediate retention of RNAs on chromatin. The broad, genome-wide binding pattern of U1 snRNA on the majority of lncRNAs, chromatin-associated mRNAs and introns is consistent with the observation that U1 abundance far exceeds that of the other snRNPs despite their equal stoichiometry in spliceosomes (Baserga and Steitz, 1993; Kaida et al., 2010). Our finding is reminiscent of U1 telescripting, an intriguing activity that is independent of U2 snRNP and splicing. U1 snRNP binds to cryptic 5’ splice sites to inhibit premature cleavage and termination at cryptic PASs scattered throughout introns, thereby promoting elongation and mRNA integrity (Berg et al., 2012; Kaida et al., 2010). Additionally, the functional interplay between U1 and PAS regulates promoter directionality and transcription elongation at transcriptional pause sites (Almada et al., 2013; Chiu et al., 2018; Core et al., 2014; Ntini et al., 2013; Richard and Manley, 2013). U1 snRNP suppresses proximal cryptic PASs enriched at transcriptional pause sites to reinforce transcription elongation in the sense direction, whereas low U1 snRNP recognition and a high density of PASs in the upstream antisense region promote early termination of PROMPTs or TSSa-RNAs. Taking our results together with these reports, we believe that U1 snRNP has additional functions beyond its well-known role in splicing.

RNA-seq analysis of small RNAs in different subcellular locations has shown that all spliceosomal snRNAs are clearly enriched on the chromatin (Tilgner et al., 2012). In addition, U1 snRNA binds to regulatory DNA elements, including the TSSs, enhancers and super-enhancers as shown by U1 RAP-DNA sequencing, probably in a base-pairing independent manner (Figure S7A) (Engreitz et al., 2014; Patel et al., 2007; Spiluttini et al., 2010). Moreover, a global RNA-chromatin interactome analysis reported that U2 snRNA binds widely across the genome on chromatin (Li et al., 2017). As splicing releases the U1 snRNP from pre-mRNAs (Will and Luhrmann, 2011), we postulate that continuous associations with U1 snRNPs without eliciting a cleavage reaction may keep U1-interacting RNAs on the chromatin. As splicing is predominantly co-transcriptional for mRNA genes (Tilgner et al., 2012), we speculate that U1-mediated chromatin tethering of RNAs could be co-transcriptional. For lncRNA-chromatin association, U1 snRNP appears to play a more dominant role than U2 snRNP. For PROMPTs and eRNAs, U1 snRNP acts synergistically with U2 to promote their chromatin association and to prevent their nuclear export. The observation that U2 snRNA also contributes in part to the chromatin association of noncoding transcripts suggests a general involvement of the splicing machinery in regulating RNA chromatin association.

Notably, the genome-wide binding of U1 snRNA is positively correlated with the level of chromatin-bound RNA, but inversely with gene expression. U1 snRNP promotes the chromatin binding of GFP-enChr (U1) RNA regardless of its expression (Figures 3D, 4C-E). On the other hand, the U1 motif decreased the transcript level of GFP-enChr RNA in the presence of a PAS but not a 3’ *Malat1* termination signal, consistent with a reported role for U1 in inhibiting gene expression in a PAS-dependent manner (Figure 3D). Interestingly, intron-retaining mRNAs, lncRNAs, PROMPTs and eRNAs share many similar features including inefficient or no splicing, chromatin association, low-level expression and short half-lives (Boutz et al., 2015; Braunschweig et al., 2014; Clark et al., 2012; Li et al., 2016). Intronic RNAs derived from both mRNA and lncRNA genes exhibit the highest levels of U1 binding (Figure S5C); yet they seldom leave the nucleus and are quickly degraded. U1 snRNP may act via the U1-PAS axis to promote RNA decay and instability. Rapid turnover renders these transcripts less likely to leave the chromatin. RNA instability and chromatin association thus appear to be intrinsically coupled for many chromatin-bound unstable transcripts. U1 snRNP-mediated RNA chromatin retention prevents nuclear export of noncoding transcripts into the cytosol from overwhelming the translation machinery (Ogami et al., 2017).

In summary, we propose that intrinsic U1 recognition sites embedded in RNA transcripts act as a key cis-regulatory signal for RNA chromatin association (Figure 7). While depleted in exons of cytoplasm-enriched mRNAs, U1 recognition sites and binding are enriched in mRNA introns and lncRNA transcripts (both exons and introns). U1 snRNPs bind to nascent lncRNA transcripts through U1:RNA base-pairing once they emerge from RNA polymerases (Figure 7A). For PROMPTs and eRNAs, U1 binding on chromatin is enriched at enhancer DNA sequences and the TSSs (the 5’ end of PROMPTs) despite the fact that U1 recognition sites are depleted in PROMPT DNA sequences. These noncoding transcripts remain associated with U1 snRNPs due to inefficient splicing or lack of splicing, and are consequently tethered on chromatin (Figure 7B). Thus, the binding of U1 snRNP is sufficient to promote RNA-chromatin association. Meanwhile, the inhibitory function of U1 snRNP on polyadenylation promotes both transcription elongation and RNA decay at cryptic and authentic PASs. Rapid RNA turnover may prevent the nuclear export of noncoding transcripts, contributing in part to their observed enrichment on chromatin (Figure 7B). Despite a more prominent requirement for U1 snRNP in regulating lncRNA-chromatin association, both U1 and U2 snRNAs are necessary to retain PROMPTs and eRNAs on chromatin. We speculate that other components of spliceosome and splicing regulators may also contribute to chromatin association of noncoding transcripts (Figure 7B). Confirmation of their involvement and of the detailed mechanisms awaits future investigation. Nevertheless, regulated chromatin retention of lncRNAs by U1 snRNP explains the remarkable phenomenon that lncRNAs as a class tend to associate on chromatin, and also provides a mechanistic insight into the functional roles of lncRNAs in modulating gene expression and chromatin structure. Elucidation of the mechanisms underlying RNA-chromatin association will facilitate an overall understanding of how noncoding portions of the genome contribute to cell-specific expression programs in development and disease.

**Figure 7.**
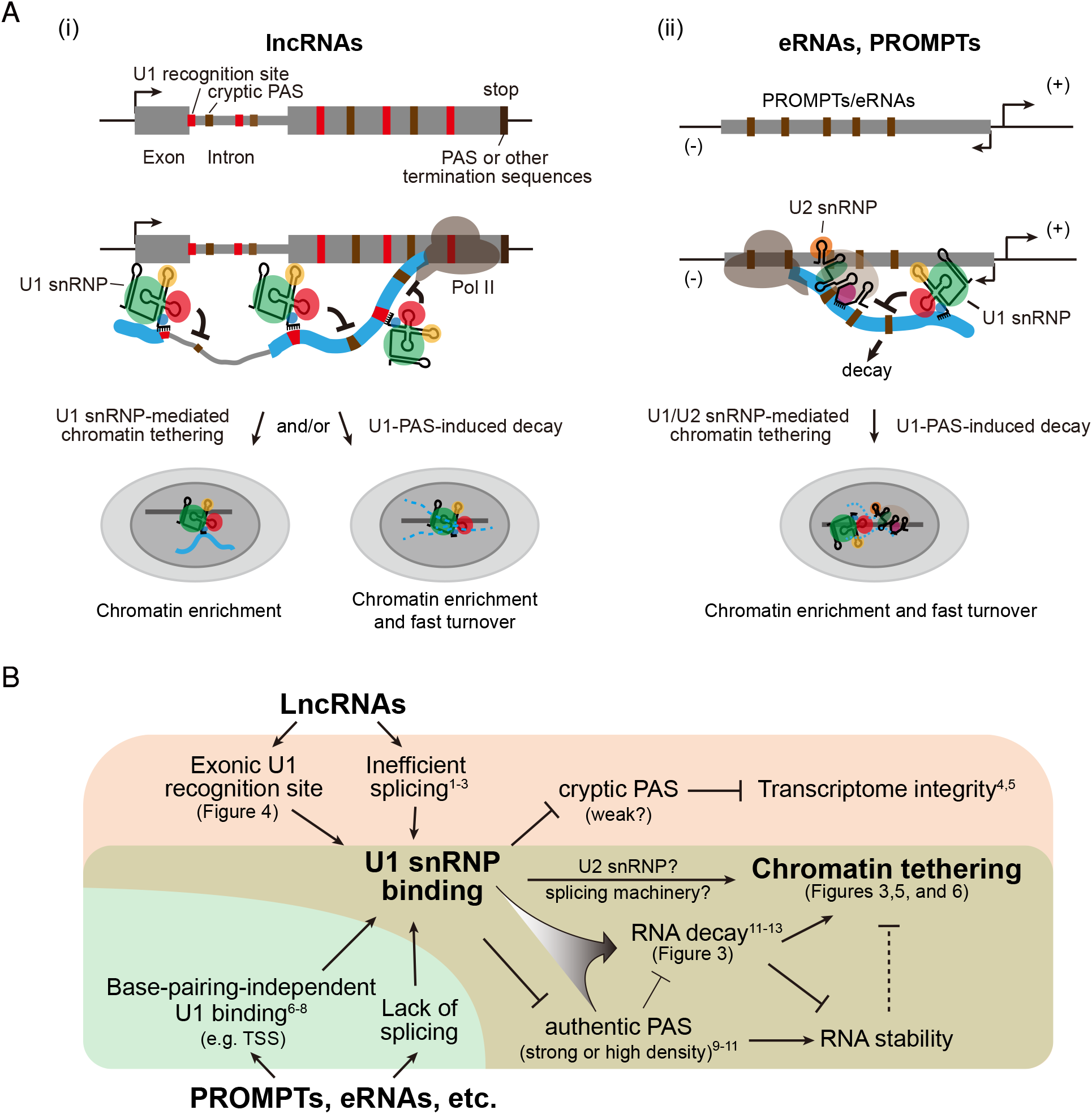
Proposed models of U1 snRNP in regulating the chromatin tethering of noncoding RNAs. (A) Models showing that the distribution of U1 recognition sites and PASs in lncRNAs (i), and PROMPTs/eRNAs (ii), contributes to U1 snRNP recognition and binding. This in turn influences the chromatin retention of lncRNAs and PROMPTs/eRNAs. For PROMPTs and eRNAs, U1 and U2 snRNPs act synergistically to promote their chromatin association. For chromatin-bound unstable lncRNAs and PROMPTs/eRNA transcripts, chromatin association and fast RNA turnover appear to be intrinsically coupled. (B) Mechanistic representation of U1 snRNP and its interplay with PAS in regulating noncoding RNA chromatin retention and decay. References shown in the model are: ^1^ Derrien et al, 2012; ^2^ Tilgner et al, 2012; ^3^ Fortes et al, 2003; ^4^ Kaida et al, 2010; ^5^ Berg et al, 2012; ^6^ Patel et al, 2007;^7^ Spiluttini et al, 2010; ^8^ Engreitz et al, 2014; ^9^ Almada et al, 2013; ^10^ Core et al, 2014; ^11^ Ntini et al, 2013; ^12^ Gunderson et al, 1998; ^13^ Goraczniak et al, 2009.

## Author contributions

Y.Y. and X.S. conceived of the project. Y.Y. performed most experiments with the help of X.Z. and Y.H., and discovered U1 motifs and performed RNA-seq analysis with bioinformatics assistance from J. L., Y.X., P. L. and Q.Z. W.S. performed total and chromatin RNA-seq. X.S. and Y.Y. wrote the manuscript.

## Acknowledgments

We thank N. Proudfoot, Z. Wang and Shen Laboratory members for insightful discussion and suggestions. Grant support is from the National Natural Science Foundation of China (31630095, 8141101062), and the National Basic Research Program of China (2017YFA0504204), and the Center for Life Sciences at Tsinghua University. Y.Y. is supported by Outstanding Postdoctoral Program of Tsinghua-Peking Joint Center for Life Sciences.

